# Sounds of the underground reflect soil biodiversity dynamics across a grassy woodland restoration chronosequence

**DOI:** 10.1101/2024.01.25.577311

**Authors:** Jake M. Robinson, Alex F. Taylor, Nicole W. Fickling, Xin Sun, Martin F. Breed

## Abstract

Fifty-nine percent of the world’s species inhabit the soil. However, soils are degrading at unprecedented rates, necessitating efficient, cost-effective, and minimally intrusive biodiversity monitoring methods to aid in their restoration. Ecoacoustics is emerging as a promising tool for detecting and monitoring soil biodiversity, recently proving effective in a temperate forest restoration context. However, understanding the efficacy of soil ecoacoustics in other ecosystems and bioregions is essential. Here, we applied ecoacoustics tools and indices (Acoustic Complexity Index, Bioacoustic Index, Normalised Difference Soundscape Index) to measure soil biodiversity in an Australian grassy woodland restoration chronosequence. We collected 240 soil acoustic samples from two cleared plots (continuously cleared through active management), two woodland restoration plots (revegetated 14-15 years ago), and two remnant vegetation plots over 5 days at Mount Bold, South Australia. We used a below-ground sampling device and sound attenuation chamber to record soil invertebrate communities, which were also manually counted. We show that acoustic complexity and diversity were significantly higher in revegetated and remnant plots than in cleared plots, both in-situ and in sound attenuation chambers. Acoustic complexity and diversity were also strongly positively associated with soil invertebrate abundance and richness, and each chronosequence age class supported distinct invertebrate communities. Our results provide support that soil ecoacoustics can effectively measure soil biodiversity in woodland restoration contexts. This technology holds promise in addressing the global need for effective soil biodiversity monitoring methods and protecting our planet’s most diverse ecosystems.

## Introduction

Soil is foundational to most terrestrial ecosystems (Thies and Grossman, 2023) and is being degraded at an unprecedented rate, largely due to unsustainable human activities (e.g., intensive agriculture, polluting by-products of industrialisation) (Ferreira et al. 2023). Soil is home to around 59% of the world’s species, making it the most biodiverse habitat on the planet (Anthony et al. 2023). Soil biota play critical roles in nutrient cycling, soil stability and structure, plant and animal health, and climate regulation (Lal et al. 2021; Banerjee and van der Heijden, 2023), among other core ecological functions. However, soil degradation poses a considerable global challenge (Wuepper et al. 2020; Prâvâlie, 2021) that needs to be ameliorated and restored to protect terrestrial biotic communities and food systems.

Biodiversity monitoring is a key part of successful restoration and there are strong calls to develop new tools and hone existing approaches to biodiversity monitoring (McGlone et al. 2020; Robinson et al. 2023a). This monitoring stage plays a crucial role in quantifying the effectiveness of interventions by measuring recovery, informing adaptive management options and potential of ongoing degradation (de Almeida et al. 2020). Ecological acoustic surveying and monitoring technology or ‘ecoacoustics’, is a rapidly growing field with efforts to expand technology to monitor more taxonomic groups. Ecoacoustics technology is used to detect the acoustic signals emitted by soniferous species, which can then be analysed to make inferences on presence/absence, abundance, composition, diversity, and distribution (Rappaport et al. 2020; Metcalf et al. 2023), as well as aspects of behavioural ecology (Abrahams and Geary, 2020). Ecoacoustics can also provide cost-effective and minimally intrusive approaches to gathering biodiversity data that can help in monitoring restoration projects (Teixeira et al. 2019; Stowell & Sueur 2020; Greenhalgh et al. 2021).

Ecoacoustics has been used to monitor elusive and protected species in several ecological contexts. For instance, passive acoustic monitoring (PAM), which involves using autonomous acoustic sensors to record biological sounds (known as “biophony”), has been widely used to detect various taxonomic groups in terrestrial ecosystems, including birds (Abrahams, 2019; Abrahams & Geary, 2020), bats (Hintze et al. 2021; López-Baucells et al. 2021), and invertebrates (Harvey et al. 2011; van der Mescht et al. 2021; Mankin et al. 2018; Mankin, 2022; Greenhalgh et al. 2023). It is also used in aquatic systems to monitor cetaceans (Jones et al. 2020; Guidi et al. 2021), amphibians (Gan et al. 2020), crustaceans (Kühn et al. 2022), and fish (Popper & Hawkins, 2019; Desjonquères et al. 2020).

Ecoacoustics has emerged as an efficient tool to measure and monitor biodiversity and has the potential to aid restoration projects (Greenhalgh et al. 2021; Znidersic & Watson, 2022). Recently, researchers have had success in applying ecoacoustics technology in the soil environment to detect and monitor soil biota. For instance, Maeder et al. (2022) demonstrated its efficacy in detecting soil meso- and macrofauna acoustic signals in silvicultural (Maeder et al. 2022) and agricultural contexts (Maeder et al. 2019). Robinson et al. (2023) showed that ecoacoustics (using specialised piezo-electric microphones) could effectively detect soil biotic communities (i.e., meso- and macrofauna) in a temperate forest restoration setting in the UK. However, there is a need to further investigate the efficacy of soil ecoacoustics in other ecosystems and biomes across the world.

Here, we apply ecoacoustics technology to measure soil biodiversity across a grassy woodland restoration chronosequence (incl. cleared, revegetated 14-15 years ago, and remnant vegetation plots) at Mount Bold, South Australia. Given that soil faunal species richness, abundance, biomass, and functional diversity have been observed to increase with time since restoration (Derhé et al. 2016), we anticipated that acoustic diversity and complexity would similarly experience an increase across our restoration chronosequence and would correlate with invertebrate abundance. Our study examined the following two hypotheses: (a) acoustic complexity and diversity will be higher in the soil recordings from remnant and revegetated plots compared with cleared plots; (b) soil acoustic complexity and diversity will positively correlate with invertebrate abundance (as per Robinson et al. 2023) and richness (as per Maeder et al. 2022), with higher scores in the remnant and revegetated plots compared with cleared plots.

## Methods

### Study location

The Mount Bold study site (35.07S, 138.42E) is a chronosequence of restoration for a *Eucalyptus leucoxylon*-dominated grassy woodland vegetation community in South Australia (Figure 1). Grassy woodland communities generally consist of a ground cover featuring grasses, herbs, and shrubs amidst sporadically distributed medium to large trees. The Mediterranean climate of the region is characterised by hot dry summers and cool winters. The site is part of a Mediterranean bioregion and is managed by South Australia’s water utility, SA Water.

**Figure 1.**
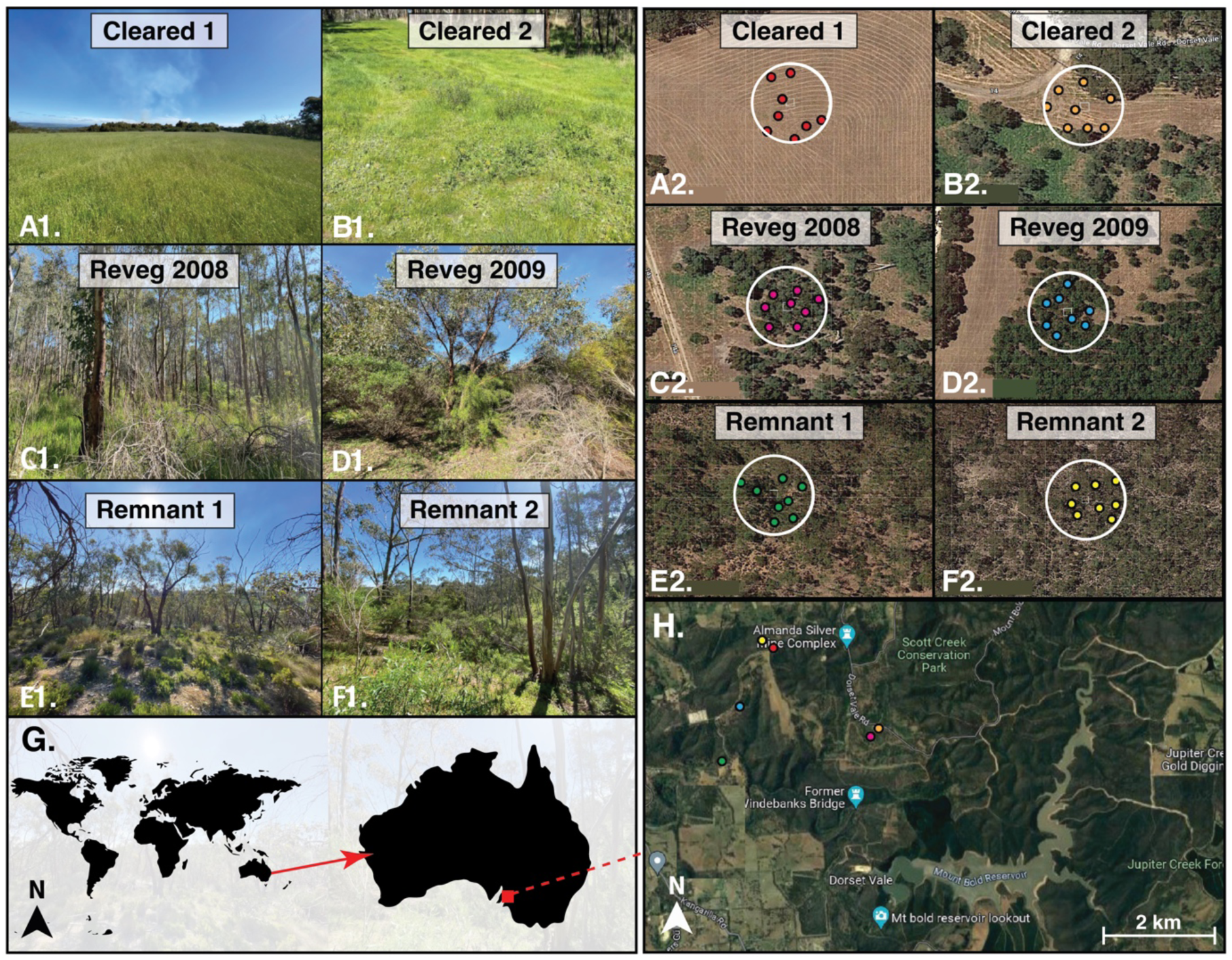
Study sites and location (Mount Bold, South Australia). (A1)-(F1): site photos for each chronosequence age class (Cleared x 2 sites, Reveg x 2 sites, Remnant x 2 sites). Sampling plots are represented by the white rings (A2)-(F2), and the 10 randomly selected sampling locations (small, coloured dots) are within each plot. (G): study location (red dot) in the broader global and Australian context, and (H): aerial view of the Mount Bold region (Basemap: Google Earth v9.175; 2023).

Mount Bold encompasses a 5,500-ha water catchment reserve located in the Mount Lofty Ranges, acknowledged as a national biodiversity hotspot (Guerin et al. 2016) and a region marked by pronounced ecological degradation (Bradshaw 2012). Before 2005, extensive portions of the Mount Bold area underwent clearance and low-density grazing for over a century. Grazing activities ceased in 2003, and a revegetation initiative was implemented from 2005 to 2009 to restore the open *E. leucoxylon* grassy woodland that characterised the catchment area before clearing (Lem et al. 2022). Consequently, a restoration chronosequence was established. The revegetation methods employed remained consistent throughout the chronosequence, involving shallow surface ripping and winter planting at all sites.

We identified three spatially independent replicate plots (at a minimum intra-treatment plot distance of 50 m apart) for sampling each chronosequence age class. Our study plots include 2 x cleared (deforested) plots (which continue to receive management in the form of slashing), 2 x revegetated plots (*E. leucoxylon* grassy woodland, revegetation commenced 14-15 years ago) and 2 x remnant plots containing relatively undisturbed *E. leucoxylon* grassy woodland communities (Fig. 1).

### Soundscape sampling

We standardised the sampling of the soundscape to occur between 10:00 and 16:00 hours in a randomly generated order (Robinson et al. 2023). We utilised a cost-effective ecoacoustics sampling device (<AU$600 for the sound recording system), with a JrF C-Series Pro contact microphone sensor (manufacturer: www.jezrileyfrench.co.uk; Yorkshire, U.K.) and a 2 m cable and a 1/4″ Neutrik jack. The C-series contact microphones offer a wide frequency response, particularly optimised for the low-end and mid-frequency range, making them well-suited for capturing soil soundscapes (Maeder et al. 2019; Gamal et al. 2020). The JrF microphone was affixed to a metal probe (a Goshawk 0.45 oz aluminium tri-peg) and connected to a handheld acoustic recording device (Zoom H4n Pro; Designed by ZOOM, Japan) before being inserted into the soil at a depth of 10 cm (Maeder et al. 2022; Robinson et al. 2023; Figure 2). We recorded Waveform Audio Format (.WAV) sound files at a 16-bit depth and a sampling rate of 48 kHz, a common rate in ecoacoustics research (Abrahams 2019), capturing sounds up to a maximum of 24 kHz, thus encompassing the entire audible range (Maeder et al. 2022). The Zoom H4n’s auto-record setting was disabled, and the gain was set to +50 dB. Each recording followed an individual probe insertion into the soil. To account for initial geophony (e.g., displaced soil particles) and potential disturbance to biophony from the physical disturbance of entering the soil, recordings consistently commenced after a preliminary 30-60 s resting period.

**Figure 2.**
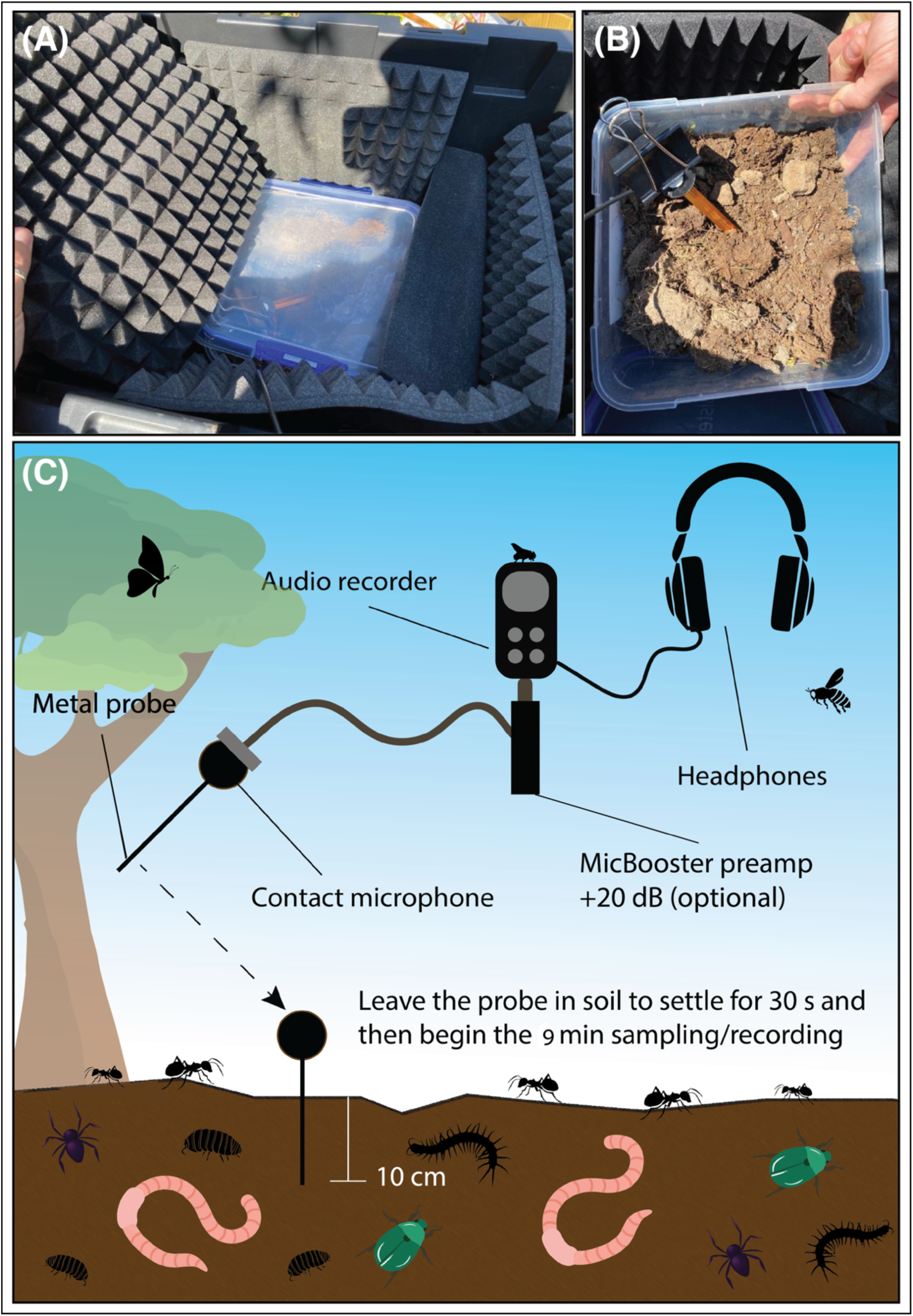
(A) The sound attenuation chamber with the foam panels, (B) the internal container that holds the soil, invertebrate samples and microphone, and (C) the in-situ study setup.

We selected soil acoustic sampling locations using a geographical information system (GIS). We created polygon boundary shapefiles around each of the six sampling plots and generated 8 random sampling points for each plot using the random points algorithm in QGIS (version 3.24.3 “Tisler”). We used one ecoacoustics system per sampling point. Soil acoustic samples were collected from the eight predetermined random points within each plot (two plots per chronosequence age class), and we recorded all eight samples simultaneously for nine minutes (a minimum of 3 minutes is recommended; Robinson et al. 2023). We repeated the sampling on five occasions in September and October 2023, and inter-site sampling was randomly selected using a digital randomiser.

### Sound Attenuation Chamber

To standardise the soil volume and, thus, acoustic detection space across treatments, we excavated soil samples. This involved collecting soil samples from eight random points in the plot with a shovel, filling a 3 L plastic container and placing this into a sound-attenuation chamber (Robinson et al. 2023). This allowed us to record a “snapshot” of the soundscape under controlled conditions (Fig. 2A-B). We used the same recording equipment for the in-situ (i.e. the source location of the soil) (Fig. 2C) and sound-attenuation chamber samples. It was important to take this sound attenuation chamber approach because recording acoustic samples in the field may capture sounds from variable detection spaces, therefore, the data acquired may not be comparable between sampling sites (Darras et al. 2016). In total, we collected *n* = 30 chamber samples (1 per plot, per visit; each sample was 9 min). The sound chamber design comprised a 60 L plastic chamber, with sound-attenuation foam installed on each internal wall, including the base and lid (Fig. 2A-B).

### Invertebrate counts

We assessed the abundance and diversity of macrofauna in the soil by utilising the soil collected during the 3 L sound attenuation chamber tests from eight random points at each plot. Following the completion of the chamber recordings, we identified and counted the invertebrates by turning over the soil from the 3 L container onto the lid of the sound attenuation chamber. We then systematically searched through the soil, adopting a left-to-right approach and carefully displacing soil particles to unveil the invertebrates, as described by Stroud (2019) and Robinson et al. (2023). On-site, we recorded the identified invertebrates in a spreadsheet, and for any invertebrates that could not be identified immediately, we conducted ex-situ identification using field photographs. Once the counting process was completed, both the soil and the invertebrates were returned to their original source location.

### Data analysis

To process the sound recordings (.WAV files), we used the wildlife sound analysis software Kaleidoscope Pro (Version 5.4.7; Wildlife Acoustics 2022). This software allows for the analysis of full-spectrum recordings to measure multiple acoustic indices, including the Acoustic Complexity Index (ACI) (Pieretti et al. 2011), Normalised Difference Soundscape Index (NDSI) (Kasten et al. 2012) and Biodiversity Index (BI) (Boelman et al. 2007) selected for this study (each described in detail below). We subsequently used these complexity/diversity (ACI and BI) and biophony-to-anthrophony ratio (NDSI) indices to test our hypotheses.

### Pre-processing steps

To provide representative sampling, reduce variability and improve the signal-to-noise ratio we pooled the eight acoustic samples collected at each plot using Audacity (v. 3.4.2) and exported them as one. WAV file per plot, per visit. Subsequently, to reduce the noise of each file and increase the signal-to-noise and the biophony-to-anthrophony ratios, we ran each file through a noise reduction pipeline using ‘pydub’ and ‘noisereduce’ libraries in Python (v 3.12). The first step in this pipeline is to load the audio files and convert them to numpy arrays. Next, noise reduction is performed, where ‘reduction_factor’ controls the amount of noise reduction, and ‘sensitivity’ adjusts the sensitivity of the noise reduction algorithm. The ‘freq_noise’ parameter controls the frequency range considered for noise reduction. In our dataset, we set this to (0, 2000) Hz to gate anthrophony while reducing noise within the biophony range.

### Acoustic Complexity Index

ACI directly measures the variability in sound intensity in both frequency and time domains, comparing the normalised absolute difference of amplitude between adjacent FFT windows in each frequency bin over seconds (K). First, ACI computes the absolute difference between adjacent values of intensity:

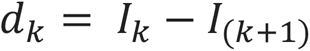

The changes in the recording’s temporal step are encompassed by the summation of the *d*′′:

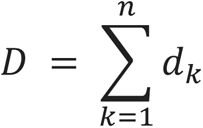

To obtain the relative intensity and reduce the influence of the distance between the microphone and biophony source, the result *D* is divided by the total sum of the intensity values (Maeder et al. 2022):

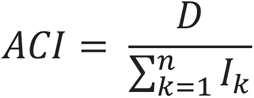

The total ACI is the sum of the ACIs across bins for each period K in the recording.

### Bioacoustic Index

BI is computed as “the area under each curve including all frequency bands associated with the dB value that was greater than the minimum dB value for each curve. The area values are thus a function of both the sound level and the number of frequency bands” (Boelman et al. 2007).

### Normalised Difference Soundscape Index

NDSI is computed as follows:

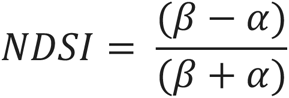

Where *β* and *α* are the total estimated power spectral density for the largest 1 kHz biophony bin and the anthrophony bin, respectively. The NDSI is a ratio in the range [− 1 to + 1], where + 1 indicates a signal containing only high-frequency biophony and no low-frequency anthrophony (Kasten et al. 2012).

As sounds above 2 kHz do not propagate well through the soil (Maeder et al. 2022), we set a maximum frequency of 2 kHz, and a lower threshold of 500 Hz for biophony in NDSI and BI in Kaleidoscope Pro.

### Statistical analysis

All statistical analysis was conducted in R Version 4.3.1 “Beagle Scouts” (R Core Team 2022) with supplementary software (e.g., Microsoft Excel for.csv file processing). To assess the distribution of the data and model fit, we applied Quantile-Quantile plots using the “qmath” function of the lattice package in R. To account for the repeated measurements within each treatment group over the five days and investigate potential correlations between invertebrate abundance and richness and acoustic index scores (ACI, BI), we applied a repeated measures correlations using the rmcorr package (Bland and Altman, 1995) in R. We also applied bootstrap resampling to assign a measure of accuracy to sample estimates for the correlations, using a minimum of 1,000 iterations. This was carried out with the boot packages in R (Canty and Ripley, 2020). To test for differences in invertebrate abundance and richness between chronosequence classes while accounting for repeated measures (plots and visits) we applied repeated measures ANOVA tests using the “aov” function in R.

We fitted linear mixed-effects models (LMEM) to the acoustic index outputs (i.e., the acoustic complexity and diverisity response variables ACI, BI) using R and its lme4 package (Bates et al. 2015). We used the Gaussian link function and the response data identity as response data were continuous. We used LMEMs because they are appropriate for including random effects of plot (i.e., each replicate of the chronosequence class) and visit (the temporal replicates, 1-5), which are essential to account for the spatial and temporal correlation between the plots and visits in our experimental design. Chronosequence class (i.e., cleared, revegetated, remnant) was included as a fixed effect. Tests of significance were conducted using Satterthwaite’s degrees of freedom t-test, which is a function of the LmerTest package in R (Kuznetsova 2020).

Soil invertebrate beta diversity was visualised using nonmetric multidimensional scaling (NMDS) ordination of Bray–Curtis distances using the Vegan package in R (Oksanen et al. 2022). The ordination plots show low-dimensional ordination space in which similar samples are plotted close together, and dissimilar samples are plotted far apart. To investigate compositional differences between treatment groups, we employed the PERMANOVA (Permutational Multivariate Analysis of Variance) using the ‘adonis’ function via the Vegan package in R (Oksanen et al. 2022). The ‘adonis’ function utilises permutation procedures to evaluate the significance of differences, providing a non-parametric and permutation-based approach. Data visualisations were produced using a combination of R and the Adobe Illustrator Creative Cloud 2022 version (Adobe, 2021).

## Results

### Soil invertebrate abundance and richness

Remnant and revegetated plots had significantly higher soil invertebrate abundance (F[1,18] = 18.99, p = <0.001; F[1,18] = 29.83, *p* = <0.001, respectively) and invertebrate species richness (F[1,18] = 25.85, p = <0.001; F[1,18] = 55.06, *p* = <0.001, respectively) than cleared plots (Fig. 3a). We noticed that soils were dominated by one particular taxon – the worker ant of *Pheidole spp*., which is <3 mm (smaller than most other species identified; Fig. 3B-E) and unlikely to create biophony within the sensitivity range of our ecoacoustics hardware. Accordingly, we ran our analysis without this taxon. The results were largely the same, with remnant and revegetated plots having significantly higher soil invertebrate abundances (F[1,18] = 27.01, p = <0.001; F[1,18] = 19.85, *p* = <0.001, respectively) and invertebrate species richness (F[1,18] = 35.52, p = <0.001; F[1,18] = 26.61, *p* = <0.001, respectively) than cleared plots.

**Figure 3.**
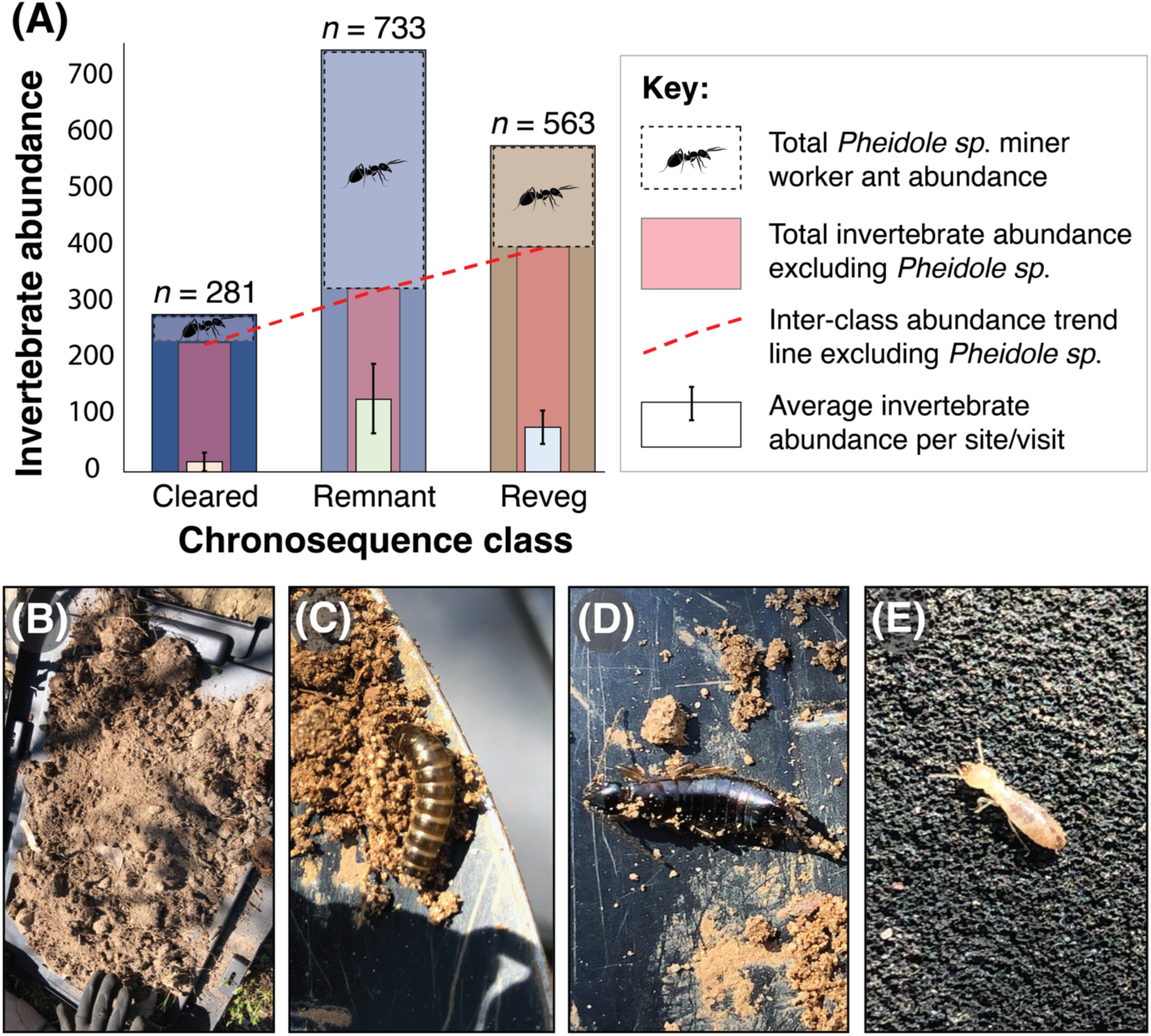
(A) Bar plots showing differences in invertebrate abundance between chronosequence classes, including post-filtering of the most abundant species (*Pheidole* sp.); (B) Soil spread on the sound attenuation chamber lid for invertebrate counting in the field; (C) unidentified beetle larvae, (D) earwig, Dermeptora, (E) termite, Isoptera.

The chronosequence class (i.e., cleared, revegetated, remnant) had a strong effect on invertebrate species richness (H = 18.9, df = 2, *p* = <0.001), with cleared having significantly lower richness compared to both remnant and revegetated treatment groups (Post-hoc Dunn tests adjusted *p* = 0.001). Remnant and revegetated treatment groups had similar richness (adjusted *p* = 0.33) (Fig. 4A-C). Visual inspection of the spectrograms for each chronosequence class also suggested a lower acoustic diversity in cleared samples (Fig. 4D). Statistical analysis of acoustic variables will follow the *Beta diversity* section below.

**Figure 4.**
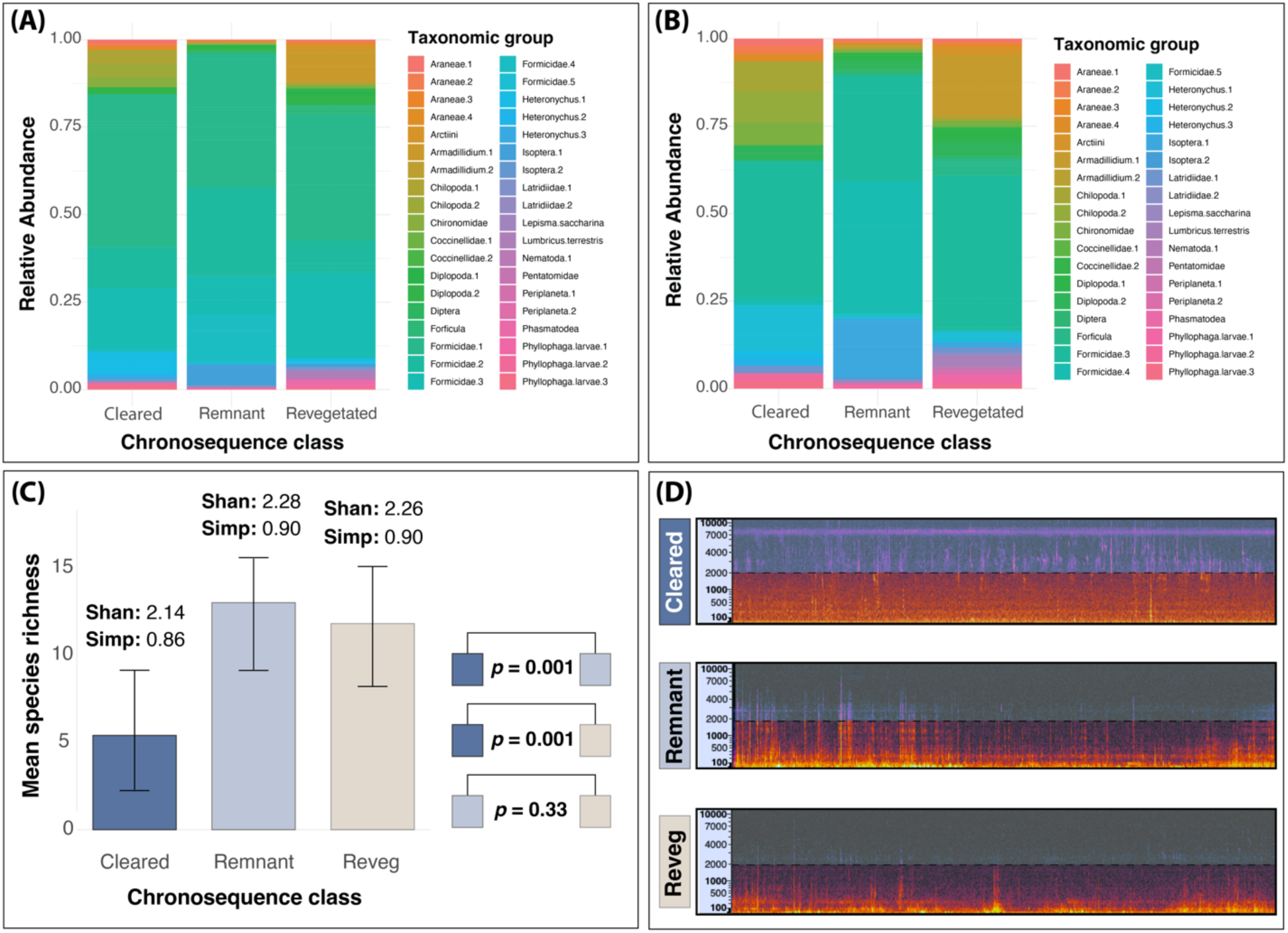
(A) Relative abundance stacked bar plots showing different invertebrate taxonomic groups between chronosequence classes; (B) Relative abundance plots showing different invertebrate taxonomic groups between chronosequence classes, but with *Pheidole spp*. filtered out; (C) Bar plots showing mean differences in species richness between chronosequence classes with corresponding Shannon and Simpson’s index scores; (D) Randomly selected spectrogram images showing a snapshot of the audio files for each chronosequence class.

### Beta diversity

Soil invertebrate community composition was significantly different between remnant and revegetated and cleared plots (stress = 0.2, r^2^ = 0.66, *p* = <0.001, permutations = 999) (Fig. 5).

**Figure 5.**
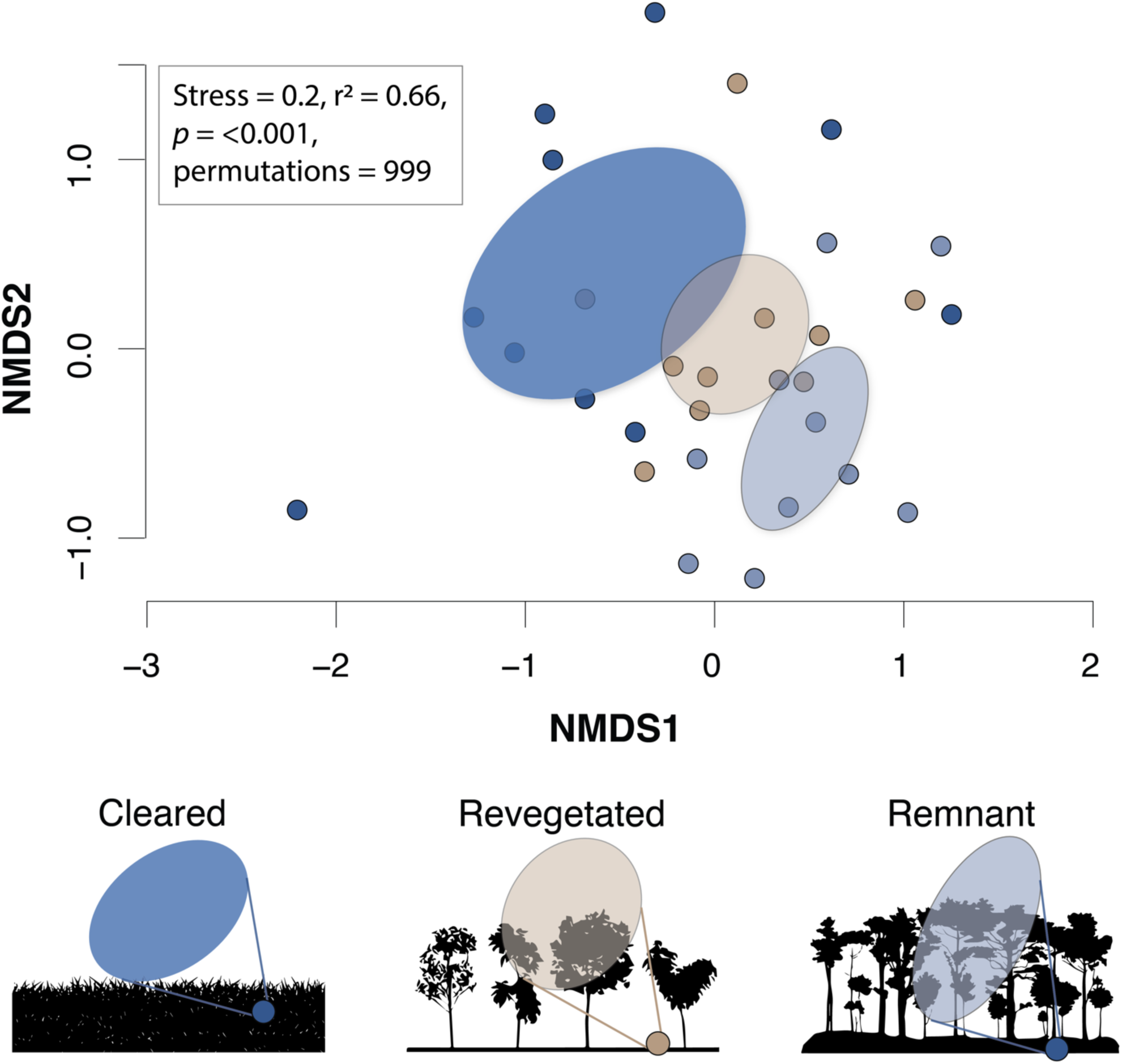
Nonmetric multidimensional scaling (NMDS) ordination plot of soil invertebrate beta diversity (community composition) for all plots (stress = 0.2; Bray– Curtis dissimilarity). Ellipses represent the confidence regions (95%) around group centroids. Clusters suggest clear differences between communities of the different treatment groups, as indicated by the coloured points and ellipses for cleared plots (dark blue), remnant plots (light blue), and revegetated plots (light brown).

### Ecoacoustics

Chronosequence class had a strong effect on ACI (Est. = 5.7, SE = 0.4, df = 28, *p* = 0.001) and BI scores (Est. = 19.6, SE = 2.6, df = 28, *p* = 0.001) in-situ (Fig. 6A-D) and on ACI (Est. = 12.4, SE = 0.55, df = 28, *p* = 0.001) and BI scores (Est. = 17.3, SE = 2.0, df = 28, *p* = 0.001) in the sound attenuation chamber (Fig. 7A-D). The acoustic index scores were higher in remnant and revegetated treatment groups compared to the cleared group.

**Figure 6.**
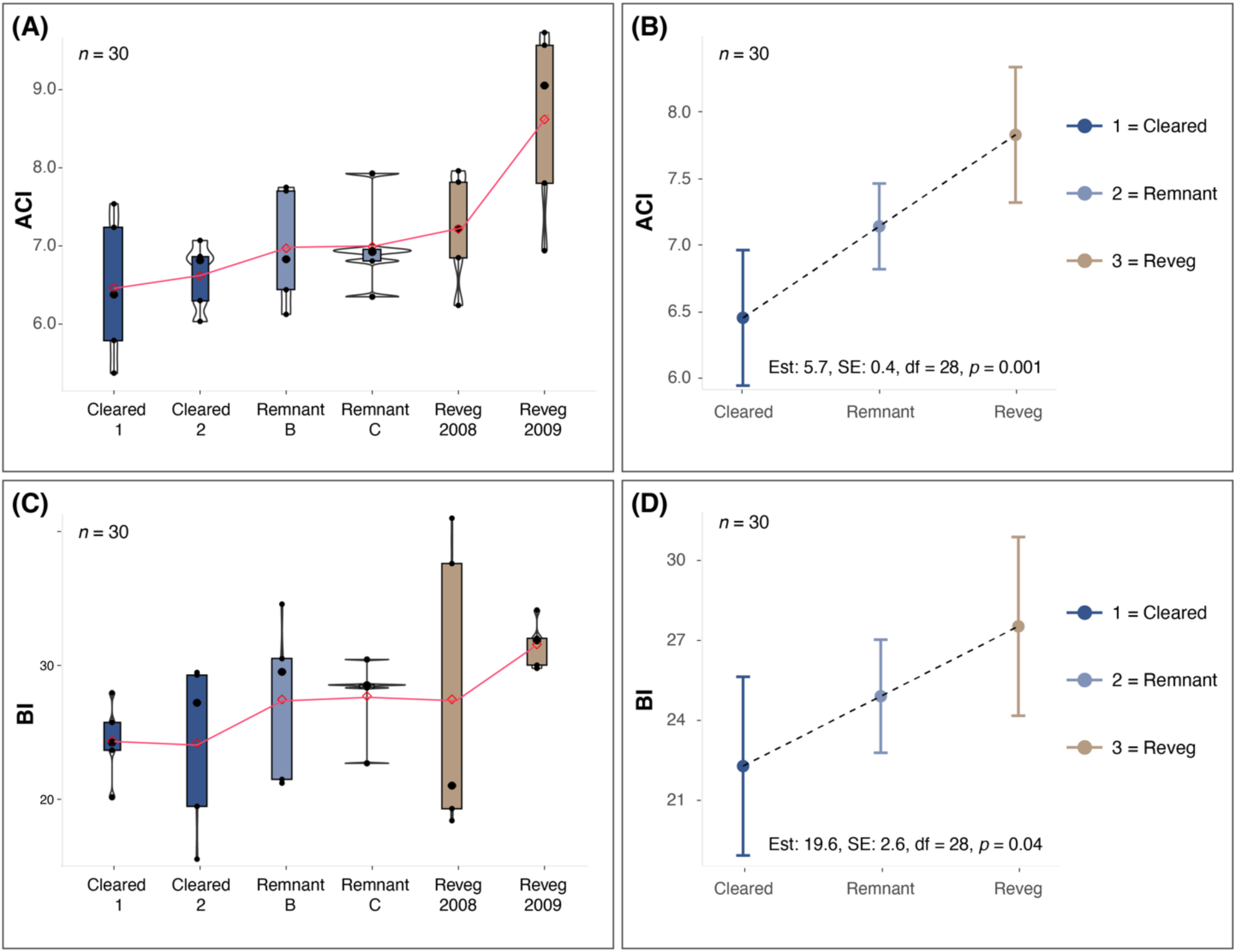
(A) Boxplot of ACI outputs for in situ (i.e. soil) samples, separated by treatment group (cleared, remnant, revegetated plots). Each plot has a red guideline to show trends in the mean values. Violins (the undulating outline around the boxplot) represent the kernel density estimation; (B) LMEM regression plot showing data observations for ACI, error bars indicating 95% confidence intervals, and dashed trend lines for each level of the factor variable; (C) Boxplot of BI outputs for in situ samples, separated by treatment group (cleared, remnant, revegetated plots); (D) LMEM regression plot for BI.

**Figure 7.**
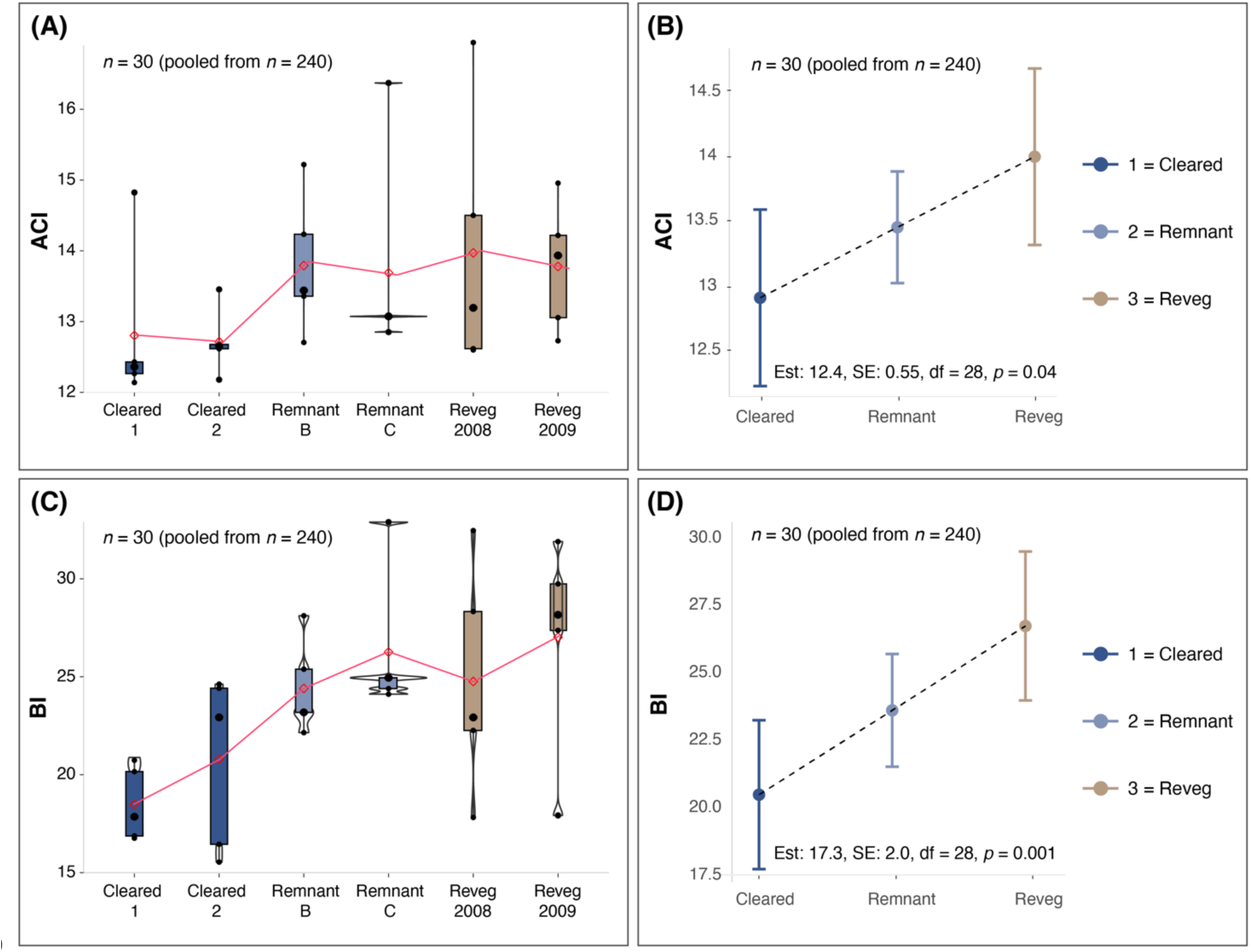
(A) Boxplot of ACI outputs for sound attenuation chamber samples, separated by treatment group (cleared, remnant, revegetated plots); (B) LMEM regression plot showing data observations for ACI; (C) Boxplot of BI outputs for sound attenuation chamber samples, separated by treatment group (cleared, remnant, revegetated plots); (D) LMEM regression plot for BI.

### Ecoacoustics and invertebrate species abundance and richness associations

Invertebrate abundance had a strong and positive effect on both ACI (‘cleared’ clusters highlighted by the light blue ellipse; Fig. 8A; β r_s_ = 0.67 (0.18, 0.90), df = 13, *p* = 0.001) and BI (β r_s_ = 0.60 (0.22, 0.83), df = 13, *p* = 0.02; Fig. 8B). Invertebrate species richness had a strong and positive effect on ACI (Fig. 8C; β r_s_ = 0.69 (0.16, 0.93), df = 13, *p* = 0.001) and a weak effect on BI (β r_s_ = 0.45 (−0.12, 0.84), df = 13, *p* = 0.09; Fig. 8D).

**Figure 8.**
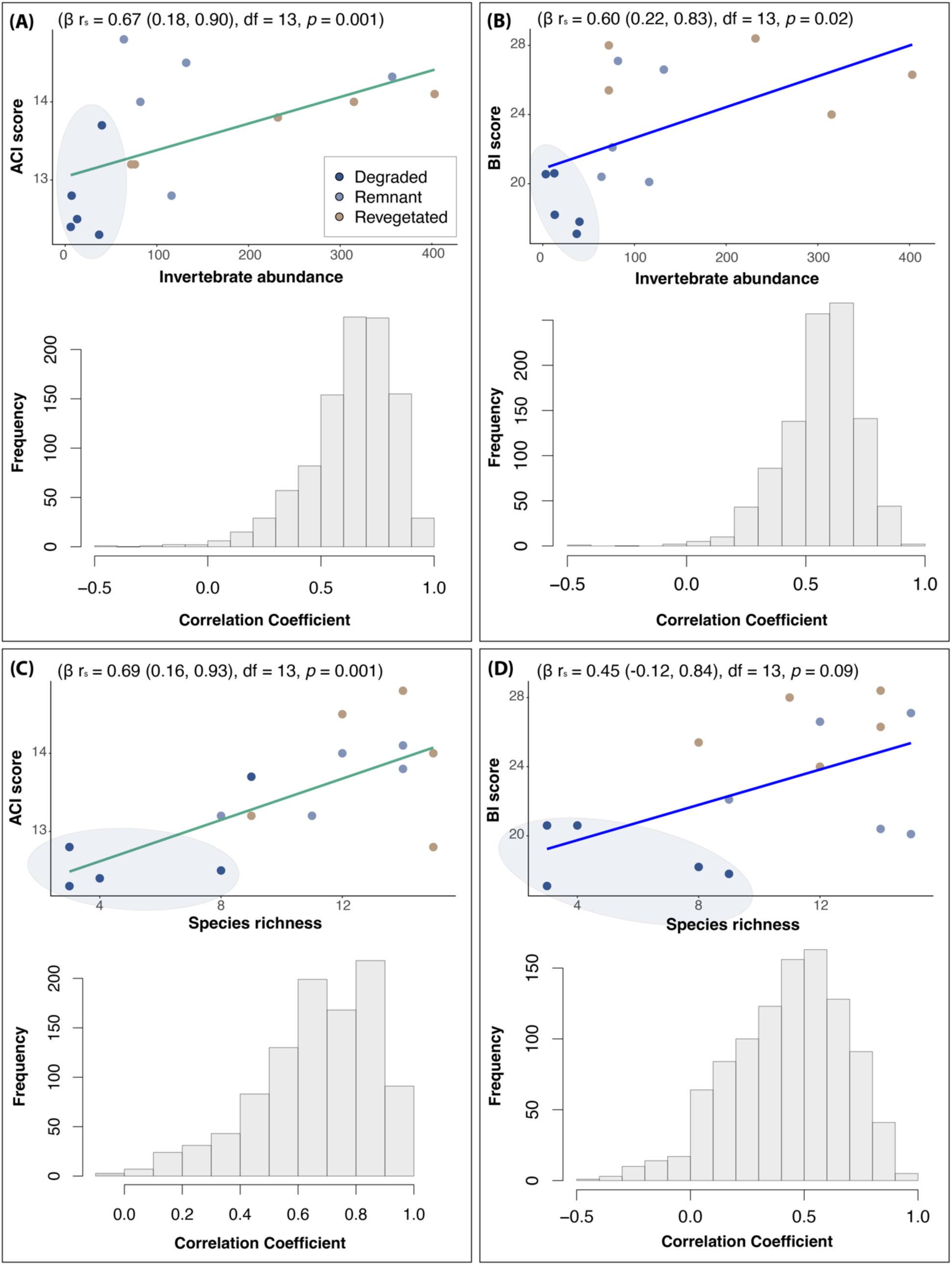
Scatter plots and bootstrap coefficient distribution plots (histograms) showing the results of the repeated measures correlations of invertebrate abundance on ACI (A) and BI (B), and invertebrate species richness on ACI (C) and BI (D). The green correlation trend line of best fit is for ACI and blue is for BI, and the data points are separated by their corresponding chronosequence class. The data points from the cleared plots are highlighted by the light blue ellipse for visual aid purposes.

## NDSI

Chronosequence class had no effect on NDSI in either the sound attenuation chamber soils (F = 0.81, df = 24, *p* = 0.55) or the in-situ soils (F = 1.85, df = 24, *p* = 0.14). All NSDI values fell between +0.94 and 0.99, indicating a very high biophony-to-anthrophony ratio for each plot (Fig. S1).

## Discussion

We show that soils in remnant and restored grassy woodland plots exhibited higher acoustic complexity and diversity than soils from cleared plots in the same area. Secondly, we demonstrate that soil acoustic complexity and diversity positively correlated with soil invertebrate abundance and richness, with higher scores in the remnant and revegetated plots compared with cleared plots. Accordingly, our study strengthens the evidence that soil ecoacoustics has the potential to measure and monitor soil biodiversity across different biomes and restoration contexts. We developed novel pre-processing steps to prepare audio files for analysis, which helps to advance the ecoacoustics field. Moreover, our beta diversity analysis suggests a strong and clear recovery in the soil biotic community, which is consistent with eDNA-based monitoring methods in the same restoration chronosequence (Yan et al. 2018).

### Soil invertebrates in the forest restoration chronosequence

Based on our invertebrate counts in the field, we observed that each chronosequence class exhibited a distinct invertebrate assemblage. Notably, the invertebrate communities in the remnant plots showed greater similarity to those in revegetated than cleared plots. Invertebrate abundance and species richness also differed significantly between the chronosequence classes, with higher counts in the remnant and revegetated plots than in the cleared plots. Although there are inconsistent global trends for the effect of restoration plantings on soil invertebrate abundance and species richness compared to control, reference ecosystems (Parkhurst et al. 2021), our results are supported by studies that show higher soil invertebrate abundance in ecosystems with lower anthropogenic disturbance (Smith et al. 2008; Nkem et al. 2020). Guarderas et al. (2022) also found that native forests exhibited greater soil invertebrate richness, evenness, and diversity than other anthropogenically altered landscapes. Moreover, our previous study (Robinson et al. 2023) showed that invertebrate abundance (but not richness) was significantly higher in restored plots compared to cleared plots. Interestingly, Robinson et al. (2023) found that earthworms were significantly more abundant in restored forest plot soils than degraded ones, which corroborated other studies (Wodika et al. 2014; Singh et al. 2020). However, earthworm abundance was very low in our South Australian grassy woodland samples, whereas ants were the most abundant taxonomic group, as found in other Australian soil invertebrate research (Nkem et al. 2020). This is likely to affect the soundscape parameters that were detected in our study.

As noted previously (Robinson et al. 2023), it might be prudent to take a more robust approach to invertebrate counting, such as using the Berlese method (Sabu & Shiju 2010). This involves specially adapted funnels to separate soil invertebrates from litter and particles and counting ex-situ (Maeder et al. 2022). That being noted, the Berlese method is more advantageous for identifying and quantifying invertebrates that are challenging to observe during field counting approaches (such as the approach taken in our study). Nonetheless, it is particularly useful for identifying smaller invertebrates that are less likely to generate a detectable biophony using our current hardware. Therefore, it would also be advantageous to improve the sensitivity and accuracy of the microphones and other audio equipment used for soil ecoacoustics (e.g., developing probes that are taxonomic group-specific). It might also be that different hardware is required to detect invertebrates that predominantly move above-ground (e.g., on the soil surface or in leaf litter using a plate-style probe design) to invertebrates that are predominantly subterranean (e.g., which the current metal peg-style probe is used for). It could also be worthwhile to explore different depths with the probes, as there is likely to be vertical stratification or zonation of soil faunal communities (Eller et al. 2018).

We show that the composition of soil invertebrate communities in revegetated plots was similar to remnant plots and very dissimilar to cleared plots. This aligns with microbial ecology research done in the same restoration chronosequence (Yan et al. 2018), which revealed a shift in the soil fungal community towards that of the natural fungal community after 10 years of active native plant revegetation. These compositional dynamics, including invertebrate abundance and richness, are reflected in our ecoacoustics analysis, as described below.

### Soil ecoacoustics in the forest restoration chronosequence

The dynamics of soil biophony using ecoacoustics have been examined in temperate (Maeder et al. 2022) and tropical ecosystems (Metcalf et al. 2024). A study conducted by Robinson et al. (2023) was the first to explore soil ecoacoustics in a forest restoration context and this was also in a temperate ecosystem, in the UK. However, our study is the first to investigate soil ecoacoustics dynamics in a southern hemisphere forest restoration context (i.e., in a South Australian grassy woodland) and the first to study soil ecoacoustics using a restoration chronosequence (space for time) framework. Specifically, we relate the acoustic complexity and diversity of soils (via the ACI and BI) to the abundance and richness of directly measured forest soil invertebrates.

We reveal significant differences in the acoustic complexity and diversity between remnant and revegetated plot soils and cleared plot soils when measured in-situ and in a sound attenuation chamber, with higher and similar scores in the remnant and revegetated plot soils. We further associate these differences with soil invertebrate abundances, which corroborates our recent study (Robinson et al. 2023). However, we also show that in contrast to Robinson et al. (2023), but in support of Maeder et al. (2022), invertebrate species richness has a strong effect on acoustic complexity (but not diversity). The correlation observed between acoustic index outputs and soil faunal communities, along with the differences noted between remnant/revegetated and cleared plots, suggests that ecological recovery of forest soils can be assessed and monitored via soil soundscapes.

Despite our results showing clear associations in both sound attenuation chamber and in-situ soils, the in-situ approach has the benefit of being less intrusive (i.e. no soil excavation is required). Therefore, future research should aim to further optimise the in-situ sampling strategy (e.g., via improved hardware and analytical pipelines) to improve the application of soil ecoacoustics in restoration and other environmental contexts.

Contrasting with previous findings (Robinson et al. 2023), there were no significant differences in NDSI values between any chronosequence class for the sound attenuation chamber soils and the in-situ soil samples. All mean NSDI values fell between +0.94 and 0.99, suggesting a consistently high biophony-to-anthrophony ratio for each plot. We previously reported a greater high-frequency to low-frequency ratio (biophony-to-anthrophony) in restored soil compared with degraded soils. This may have been due to the greater earthworm activity changing the temperate soil characteristics (making them more air permeable) to allow better propagation of higher-frequency sounds, thereby increasing NDSI values (Keen et al. 2022). However, we view our very high biophony-to-anthrophony ratio result as a positive facet as it suggests that disturbance via anthrophony was very low in our samples.

Inter-ecosystem variability and the variety of soil acoustic signals made by fauna are still poorly understood, as are diel elements, in which some soil organisms are more active and therefore potentially producing louder and more diverse sounds at certain times of the day than others. This temporal element was not captured by our study and warrants further research.

## Conclusion

Our study provides valuable insights into the use of soil ecoacoustics as a non-invasive tool for assessing soil biodiversity in forest restoration contexts. We confirm that soils in both remnant and revegetated South Australian grassy woodland plots exhibit higher acoustic complexity and diversity compared to cleared plots, which associated with invertebrate abundance and richness in these soils. Our work is pioneering by investigating soil ecoacoustics within the restoration chronosequence framework and showcasing its potential as an important tool for assessing soil health. We emphasise the need for further optimisation of in-situ sampling strategies and consideration of temporal (and soil depth) dynamics in future research to enhance the applicability of soil ecoacoustics in biodiversity monitoring. This study, along with others in the growing field of soil ecoacoustics, offers a novel and promising avenue for understanding soil biodiversity in different ecosystems and restoration contexts.

## Acknowledgements

XS acknowledges funds from the National Natural Science Foundation of China (No. 32361143523), and the International Partnership Program of the Chinese Academy of Sciences (No. 322GJHZ2022028FN).

## Supplementary materials

**Figure S1.**
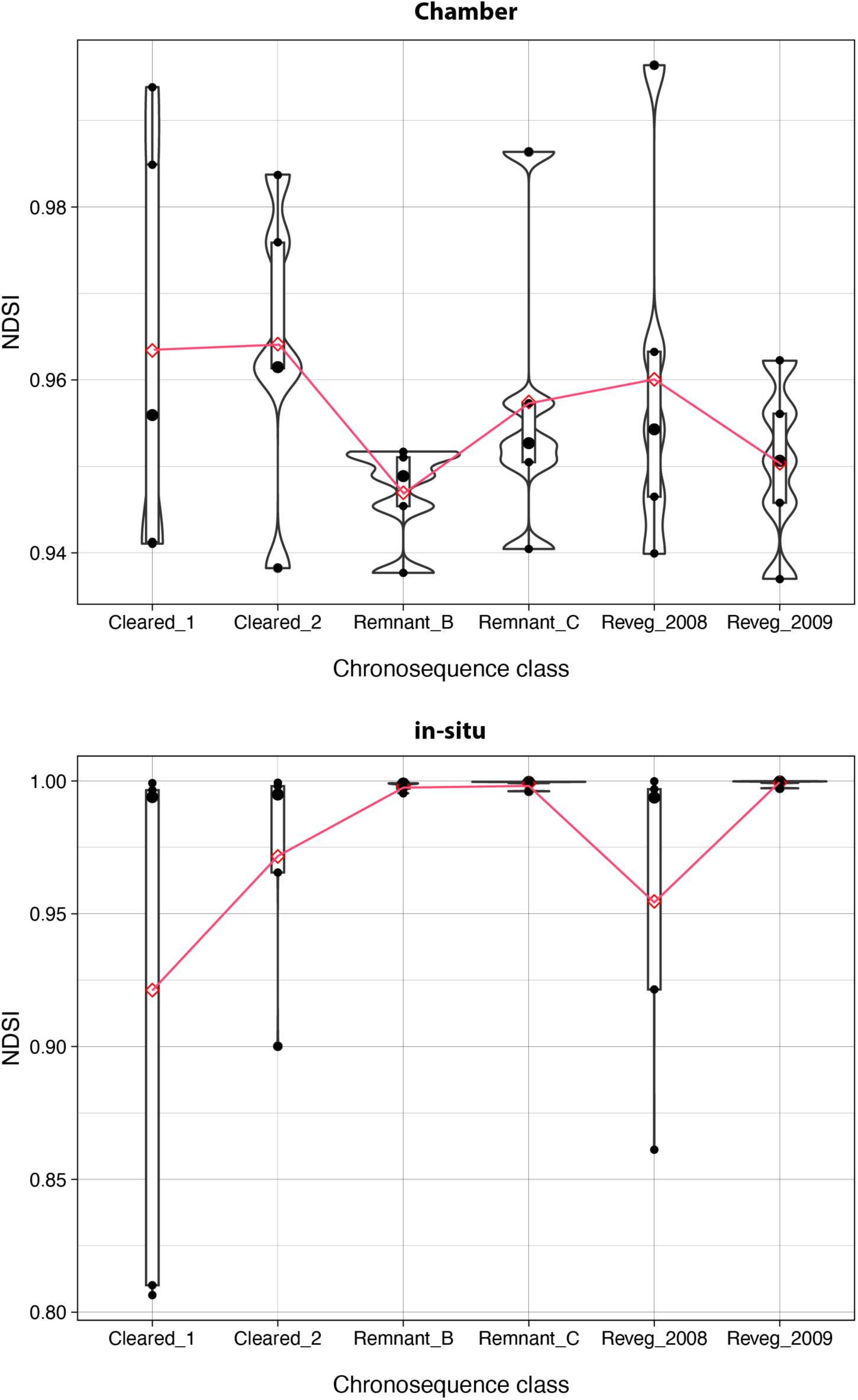
Boxplots showing NDSI results for each chronosequence class in both sound attenuation chamber soils (top panel) and in-situ soils (bottom panel).

