## Supplementary Figure S1 for "Sounds of the underground reflect soil biodiversity dynamics across a grassy woodland restoration chronosequence"

### Supplementary materials

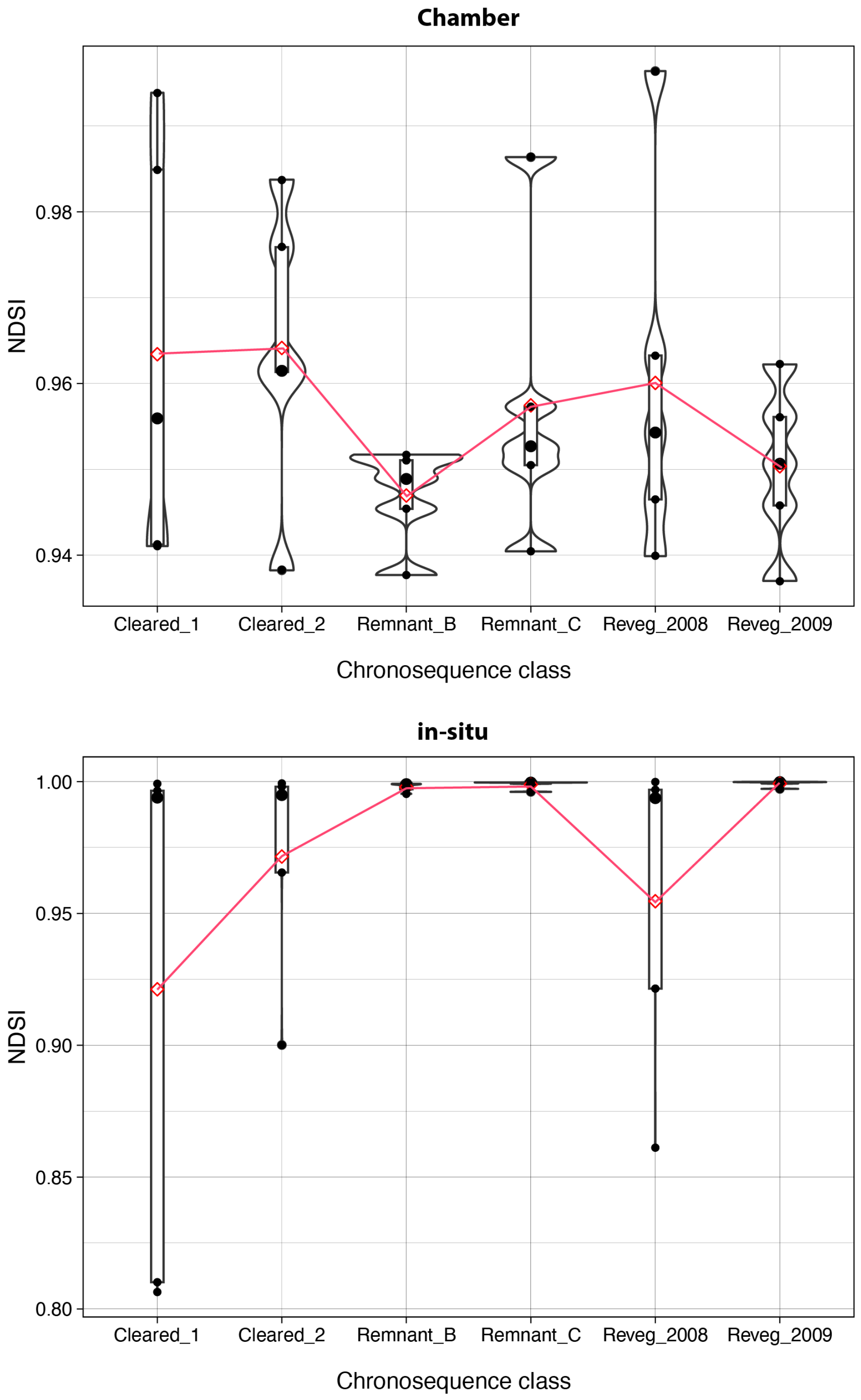


**Figure S1 |** Boxplots showing NDSI results for each chronosequence class in both sound attenuation chamber soils (top panel) and in-situ soils (bottom panel).
